# Super-resolution microscopy reveals the nanoscale organization of the DEK protein cancer biomarker

**DOI:** 10.1101/2023.02.28.530467

**Authors:** Agnieszka Pierzynska-Mach, Alberto Diaspro, Francesca Cella Zanacchi

## Abstract

The chromatin architectural factor DEK is involved in regulating the chromatin organization and is highly overexpressed in numerous forms of cancer. DEK is traditionally described as uniformly distributed within the nucleus by diffraction-limited microscopy studies, but super-resolution advent highlighted the formation of cluster-like DEK structures. Still, a characterization of the DEK protein cellular distribution and its role in cancer and cell proliferation is missing. In this work, we used single-molecule localization microscopy (SMLM) to investigate the nanoscale characteristics of DEK organization in normal-like and aggressive types of breast cancer cell lines, characterizing the number of localizations per cluster, as well as areas and density of clusters. We show how the cluster features correlate with the breast cell type and how the chromatin decompaction influences the DEK clusters in both cell lines. Our results suggest that the DEK density and nano-organization are preserved and are not influenced by protein overexpression itself or by chromatin compaction changes.

## 1. Introduction

Cancer is one of the leading causes of mortality and can originate from the alteration of genetic material or by heritable features of epigenetic characteristics (1). Two critical features in cancer progression are an abnormal chromatin local organization and the perturbation of the nanoscale landscape of the nuclear factors (2,3).

One of the non-histone chromatin architectural factors is DEK, which has a pleiotropic mode of action by influencing several regulatory cellular pathways. DEK is involved in chromatin structure and chromatin-related processes regulation, such as transcriptional activation and repression (4–6), DNA replication (7–10), and DNA damage response and repair (11–15). DEK binds to many highly and commonly expressed genes (5) and is involved in gene regulation in breast cancer cells (16). DEK can be observed mainly inside the cell nucleus of eukaryotic cells (4,17,18), and several studies demonstrated that it associates with open chromatin, rich in euchromatin histone marks (5,18–20). However, DEK has also been implicated in the maintenance of transcriptionally inactive heterochromatin (21,22).

Interestingly, DEK is highly overexpressed in numerous forms of cancer (23–29) and, in general, its expression is considered an indicator of the cell proliferation level (30). In the context of breast cancer, for example, Western Blot and immunohistochemistry assays showed that DEK is highly expressed in human breast cell lines (e.g., different MDAMB cell lines), less expressed in a non-tumorigenic immortalized cell line (MCF10A), and expressed the least in normal tissues (28). These assays provide critical information on the overall protein expression but clearly cannot report the effects of such overexpression on the precise spatial distribution of the protein in the cell. When observed with diffraction-limited fluorescence microscopy, DEK is commonly described as uniformly distributed within the cell nucleus (18,31). However, our recent study employing SMLM suggests the formation of cluster-like DEK structures (10).

In this study, we perform a thorough characterization of such DEK clusters for normal-like (MCF10A) and metastatic (MDAMB231) breast cancer cell lines, known to exhibit a different level of overall DEK expression, through a combination of single-molecule localization microscopy and clustering analysis approaches, aiming to uncover the topological landscape of this cancer biomarker.

## 2. Materials and Methods

### 2.1. Cell culture

Mammary epithelial, non-transformed MCF10A cells (ATCC CRL-10317) were grown in DMEM:F-12 (Dulbecco’s Modified Eagle Medium : Nutrient Mixture F-12) (1 : 1) medium (Gibco, 11330057) supplemented with 5% Horse Serum (HS), 2 mM L-glutamine and 1% penicillin/streptomycin (Sigma-Aldrich, G6784), 10 µg/ml insulin (Sigma-Aldrich, I9278) and 0,5 µg/ml hydrocortisone (Sigma-Aldrich, H0888). Human Epidermal Growth Factor (hEGF, Sigma-Aldrich, E9644) was added freshly before the use of full medium (20 nm/ml). MDAMB231 cells (ATTC HTB-26) were grown in DMEM medium (Gibco, 11330057) supplemented with 10% Fetal Bovine Serum (FBS, Euroclone, ECS0180L), 1% penicillin/streptomycin (Sigma-Aldrich, G6784) and 2 mM L-glutamine. Cells were grown on 10cm^2^ Petri Dish for a month (about 10-12 cell passages) at 37°C in 5% CO_2_. For any experiments on fixed cells, cells were plated on glass coverslips coated with 0,5% (w/v) pork gelatin (Sigma-Aldrich, G2500) previously dissolved in phosphate buffer saline (PBS) and autoclaved. For hyperacetylation experiments, MCF10A and MDAMB231 cells were treated with 300 nM of trichostatin A (TSA) (Sigma-Aldrich, T8552) in complete growth medium for 24h prior the fixation.

### 2.2. Immunolabelling and DNA staining

For the detection of the protein of interest, the cell cultures were washed with pre-warmed PBS 3x, and fixed with formaldehyde (3,7%, methanol free). After the subsequent blocking with the blocking buffer composed of 0,1% Triton X-100 and 3% w/v of bovine serum albumin (BSA) for 1h at room temperature, samples were incubated with the primary antibody (overnight at +4°C) α-DEK (mouse, Santa Cruz, sc-136222, 1:50). Secondary antibodies staining was performed with the use of F(ab’)2-Goat anti-Mouse IgG (H+L) Cross-Adsorbed Secondary Antibody, Alexa Fluor 647 (ThermoFisher Scientific, A21237, 1:250, 45min in RT). After finishing the immunostaining, samples were post-fixed in PFA 2% for 5 min and stored in PBS at +4°C. For the experiments in which the DNA in cells was stained, the cells were incubated with the DNA precursor, 5-ethynyl-2’-deoxyuridine (EdU), with the final concentration of 10 µM in complete medium, 24h prior to the fixation.

### 2.3. STORM Imaging and Data Reconstruction

#### 2.3.1. STORM microscope

A commercial N-STORM TIRF Eclipse Ti2 microscope equipped with an oil immersion objective (CFI SR HP Apochromat TIRF 100XC Oil, NA 1.49) was used to acquire 20000 frames at the frame rate 30ms per frame using oblique incidence excitation.

#### 2.3.2. STORM imaging buffer

All samples were imaged in the previously described imaging buffer with the use of GLOX component (32). The buffer contains a glucose oxidase solution as the oxygen scavenging system: 40 mg/ml catalase (Sigma-Aldrich, #C40-100MG), 0.5 mg/ml glucose oxidase (Sigma-Aldrich, G2133-250KU); and MEA 10 mM: cysteamine (Sigma-Aldrich, 30070-50G) in 360 mM Tris-HCl.

#### 2.3.3. Imaging protocol

Imaging was performed by the acquisition of 20000 frames of 647 channel with an exposure of 30 ms. For the imaging of Alexa Fluor 647 the 647 nm laser was used in order to excite and switch in to the dark state the dye. The 405 nm laser light was used in order to reactivate the dye into a fluorescent state. The imaging was performed with a repeating cycle of 1 activation frame followed by 3 read-out frames, with the use of sCMOS camera (ORCA-Flash4.0, Hamamatsu Photonics K.K.). All the measurements were performed following the same imaging protocol (supplementary data). The Nikon Perfect Focus System was applied during the entire recording process. The acquisition was performed with the use of Continuous STORM filter.

#### 2.3.4. Image reconstruction and clustering analysis

The images were reconstructed using a custom software (Insight3, kindly provided by Dr. Bo Huang of the University of California) by Gaussian fitting of the single-molecule raw images in each frame to determine the x–y coordinates. First, molecules are identified by setting a common threshold of 600 counts/pixel. Gaussian fitting provides the positions of each identified molecule and the final images were obtained by plotting each identified molecule as a Gaussian spot. Images were corrected for drift by cross correlating images obtained from subsets of frames as described in the literature (33). Molecules exhibiting a localization precision below 10nm are excluded to avoid aspecific signal and false localizations.

Cluster analysis of STORM datasets was performed with a custom-written code (MATLAB, The MathWorks, Natick, MA) implementing a distance-based clustering algorithm previously published (34,35). This density-based clustering algorithm is used to identify spatial localizations clusters, with a minimum number of localizations per cluster equal to 20 N_Loc_/cluster to select the DEK contribution and avoid aspecific background and aggregates. Briefly, a density map is obtained and transformed into binary images. From the binary images, only localizations lying on adjacent (six-connected neighbors) nonzero pixels of the binary image were considered. Initialization values for the number of clusters and the relative centroid coordinates were obtained the density map identifying local maxima within the connected region, and localizations were associated with clusters based on their distance from cluster centroids. New cluster centroid coordinates were iteratively calculated as the average of localization coordinates belonging to the same cluster. The procedure was iterated until convergence of the sum of the squared distances between localizations and the associated cluster. Selected parameters were kept constant to ensure proper comparison among the different samples.

### 2.4. Statistical Analysis

Statistical tests were performed in OriginPro, Version 2020 (OriginLab Corporation, Northampton, MA, USA) and GraphPad Prism version 9.4.1 (GraphPad Software, San Diego, California USA, www.graphpad.com). Data points corresponding to the number of localizations, cluster areas, and densities were tested with a non-parametric two-sample, two-tailed Kolmogorov-Smirnov test. In order to obtain the statistical significance level of the differences between the distributions composed of a large set of data (between 22 000 and 47 000 clusters identified), bootstrapping approach was applied. In brief, 1000 data points were randomly sampled out of each of datasets (number of localizations, areas and densities), the maximum difference was obtained by K-S test, and the process was repeated 1000 times. In that way, a mean difference with the standard deviation was obtained, which has been compared to a p-value. As a control, the data coming from the same distribution (e.g., MCF10A control sample) was sampled as above, resulting in high p-values for intrinsic data. The comparison of the distributions between samples lead to high differences and thus low p-values (not significant, n.s. when p > 0.05; significant when “ * “ meaning p ≤ 0.05; “ ** “ meaning p ≤ 0.01; “ *** “ meaning p ≤ 0.001).

## 3. Results

### 3.1. DEK distribution into nanoclusters differs between normal-like and metastatic breast cancer cells

In order to characterize DEK protein cellular distribution, we used single-molecule localization based super-resolution microscopy. This approach allowed us to quantitatively compare the DEK pattern in MCF10A and MDAMB231 cells, representing the normal-like breast basal epithelial cells and triple negative, metastatic basal breast cancer cells, respectively. Previous studies indicate that DEK protein locates mostly within the cell nucleus (4,10,18), and, more precisely, it can be associated with euchromatin (i.e., the active compartments of chromatin) (19). Considering the overexpression of DEK in breast cancer samples, we speculated that its nanoscale organization would differ if compared to normal-like cells. To verify this, we studied its cluster organization (Fig. 1). MCF10A and MDAMB231 cells were fixed and immunostained against DEK following the protocol suitable for fluorescence super-resolution microscopy (Materials and Methods). We used stochastic optical reconstruction microscopy (STORM) and clustering analysis to identify DEK protein clusters and to characterize the number of localizations per cluster, as well as areas of individual clusters. Wide-field images (Fig. 1. A, D) did not reveal major visual differences in the DEK pattern between the cell lines, however the clustering analysis gave us precise insight into the features of individual DEK clusters. In order to perform the analysis, clusters of DEK were segmented using a previously developed distance-based algorithm (34) (Fig. 1 C, F). Based on this operation, we obtained the distributions of the number of localizations per cluster (Fig. 1 G) and of the cluster area (Fig. 1 H) for each cell type. The average number of localizations per DEK cluster in MDAMB231 cells (30.7 ± 21.9) resulted in about 13% higher than in MCF10A (27.2 ± 18.7) (Suppl. Fig. 1 A, C), and the difference of the distributions was statistically significant (p ≤ 0.05). Furthermore, we observed a higher mean area of DEK clusters in MDAMB231 cells compared to MCF10A cells (about 2.6% higher) (Suppl. Fig. 1 B, D), although the difference was not statistically significant (p=0.0796). Next, we computed the distributions of the DEK clusters densities in both cell lines and observed that they exhibited no statistically significant differences (Fig. 1 I). This result brought us to hypothesize that DEK protein organizes in clusters of different protein copies across different cancer cell states. Still the density is conserved, regardless of the DEK expression level.

**Fig. 1.**
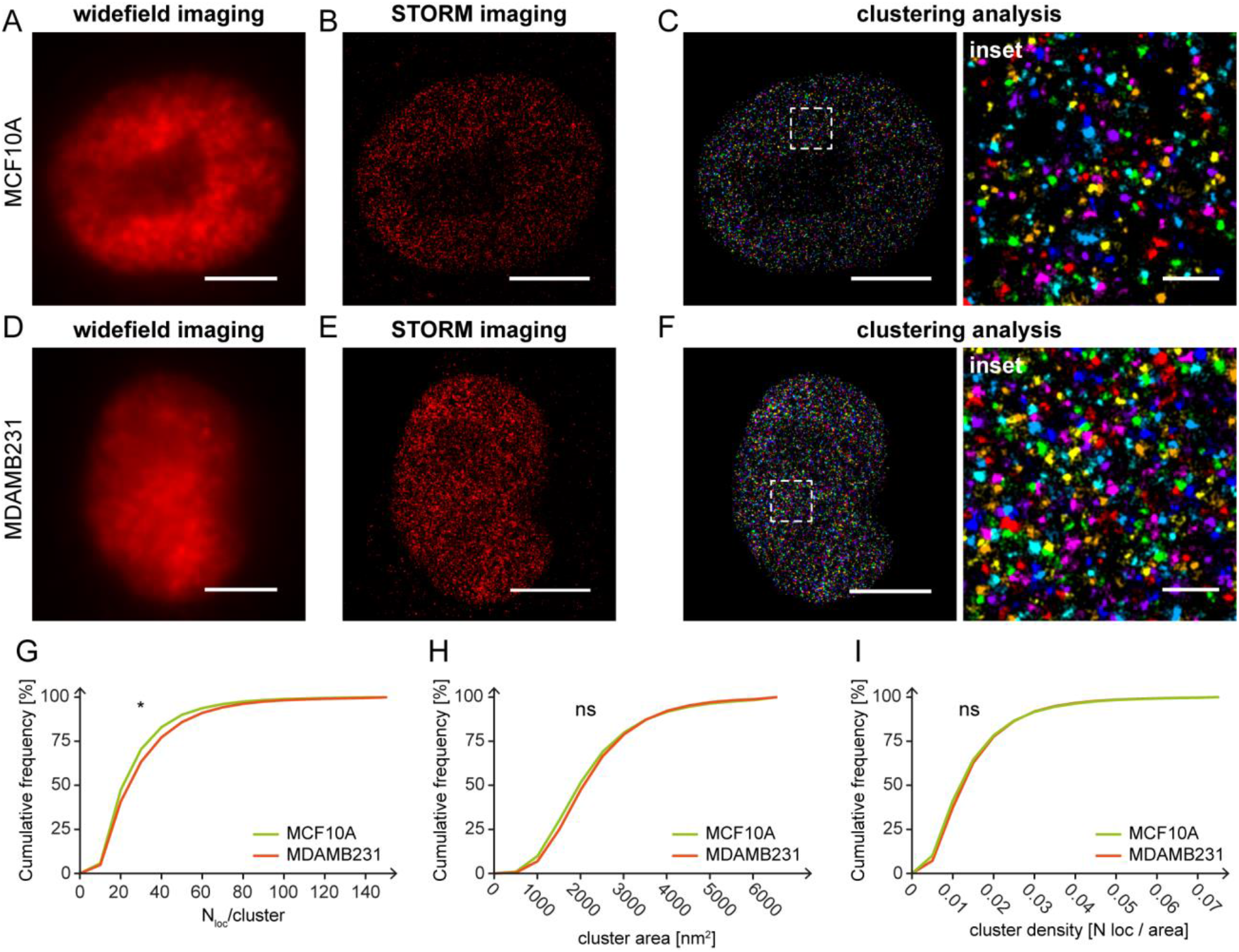
Super-resolution imaging of DEK protein clusters in breast cancer model cell lines. (A) Widefield, (B) STORM images and (C) clustering image of DEK protein in MCF10A nucleus immunolabelled with AF647 (each color represents different cluster ID). (D) Widefield, (E) STORM images and (F) clustering image of DEK protein in MDAMB231 nucleus immunolabelled with AF647 (each color represents different cluster ID). Cumulative distributions of (G) the number of localizations / cluster, (H) cluster area and (I) cluster density show the DEK cluster features differences between MCF10A and MDAMB231 cells. The average number of localized events in MCF10A cells: 27.2 ± 18.7 compared to MDAMB231 cells: 30.7 ± 21.9. The average cluster areas MCF10A cells: 2224.9 ± 1252.8 [nm^2^] compared to MDAMB231 cells: 2282.8 ± 1131.1 [nm^2^]. The average cluster densities in MCF10A cells: 0.0147 ± 0.011 [N_loc_/cluster] and MDAMB231 cells: 0.0151 ± 0.0106 [N_loc_/cluster]. The total number of analyzed clusters is: N_MCF10A_ = 26906 and N_MDAMB231_ = 22008, for n ≥ 4 independent experiments. Kolmogorov-Smirnov statistical test: not significant, n.s. when p > 0.05; “ * “ when p ≤ 0.05. Scale bar = 5 µm. Scale bar insets = 500 nm.

To investigate if this behavior is a more general trait, we decided to compare the DEK distribution between two different chromatin compaction states. Indeed, it is known that chromatin compaction alters the nanoscale distribution of chromatin components and chromatin-associated proteins (34).

### 3.2. The influence of TSA on chromatin decondensation in normal-like and metastatic breast cancer cells

In order to control the chromatin compaction, we used Trichostatin A (TSA), a well know histone deacetylase inhibitor that induces the chromatin opening (36) and leads to chromatin decompaction by global histone hyperacetylation. On the cellular level, TSA influences chromatin organization by opening the DNA due to decreased nucleosomal occupancy, which has been revealed with single-molecule localization microscopy (37).

To confirm the TSA influence on chromatin compaction in the selected cell lines, we observed the DNA density both using wide-field and SMLM imaging in control and TSA-treated cells (Fig. 2). The control samples of both MCF10A (Fig. 2 A) and MDAMB231 (Fig. 2 B) exhibit typical DNA distribution within the cell nucleus with higher fluorescence intensity coming from dense, heterochromatin regions (nuclear periphery and surrounding of the nucleolus). Lower density euchromatin regions exhibit local variations in DNA density (Fig. 2 A, B upper row, insets). The TSA treatment of MCF10A and MDAMB231 cells resulted in the global hyperacetylation of histone tails, which manifested with the genome-wide decompaction of DNA when observed with SMLM (Fig. 2 A, B lower row, insets).

**Fig. 2.**
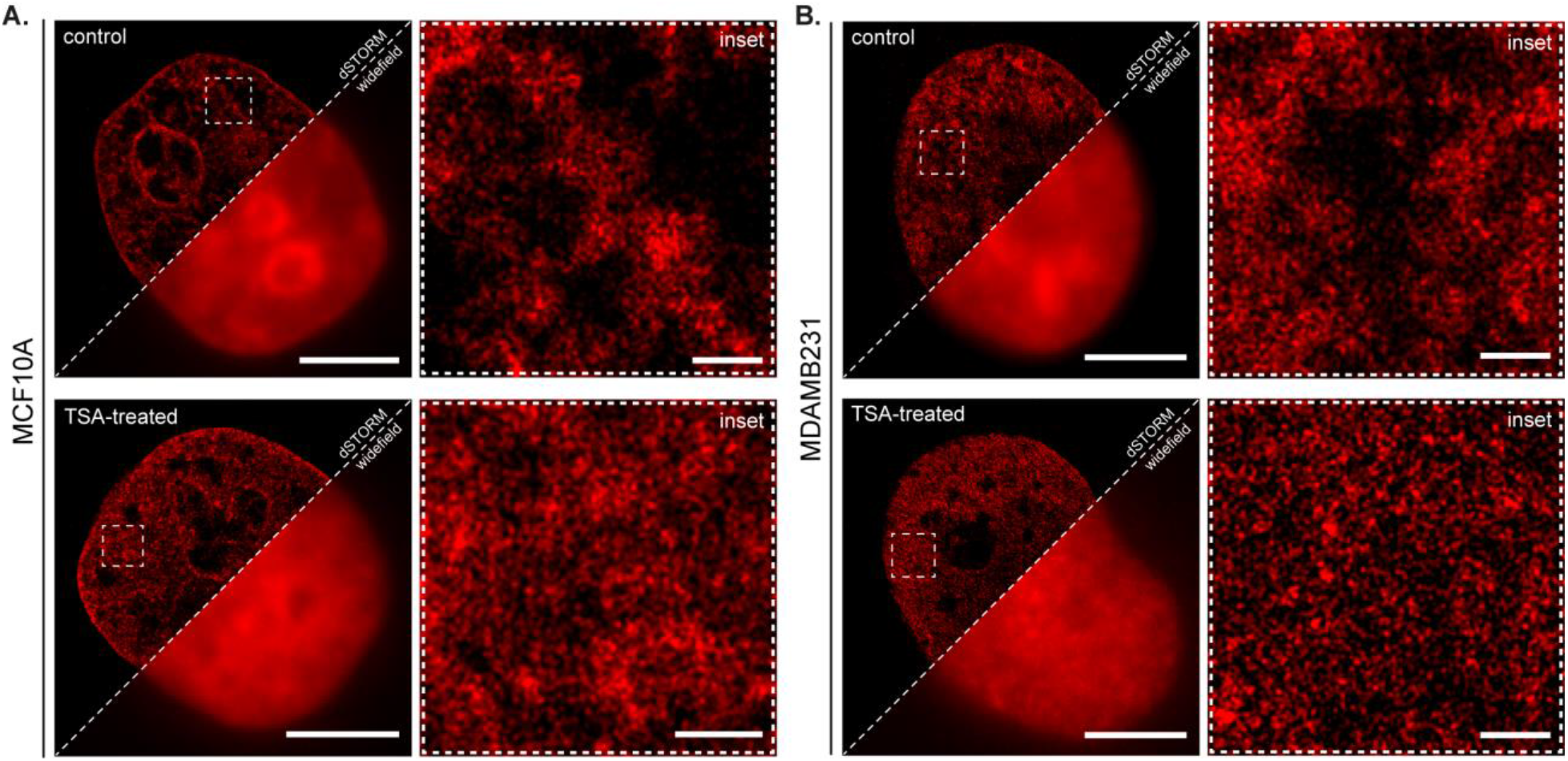
SMLM imaging reveals the influence of TSA on the DNA compaction. Widefield and super-resolution images of DNA in the nucleus labeled with EdU (incubation 24h) in MCF10A (A) and MDAMB231 cells (B); Scale: 5 µm. The insets reflect the euchromatin region within the squares in full size image. Scale: 400 nm.

These results are in accordance with previously done studies with the use of single-molecule localization microscopy, which have shown the DNA density change upon the influence of TSA in human BJ fibroblasts (37).

Once confirmed that TSA effectively inflicts chromatin decompaction, we moved to investigate the potential nanoscale redistribution of DEK clusters following the TSA treatment.

Using SMLM imaging combined with clustering analysis, we performed the analysis of the features of the DEK clusters corresponding to control cells (Fig. 3 A-C and Fig. 4 A-C) and TSA-treated cells (Fig. 3 D-F and Fig. 4 D-F). The clustering analysis provided the distributions of a number of localizations per cluster (Fig. 3 G and Fig. 4 G) and the cluster areas (Fig. 3 H and Fig. 4 H). We observed a statistically significant increase in the number of localizations per cluster in cells treated with TSA in comparison to control cells (25% and 22.1% for MCF10A and MDAMB231 cells, respectively) together with a statistically significant increase in the mean cluster area (14.3% and 17.7% for MCF10A and MDAMB231 cells, respectively) (Suppl. Fig. 1 E-H).

**Fig. 3.**
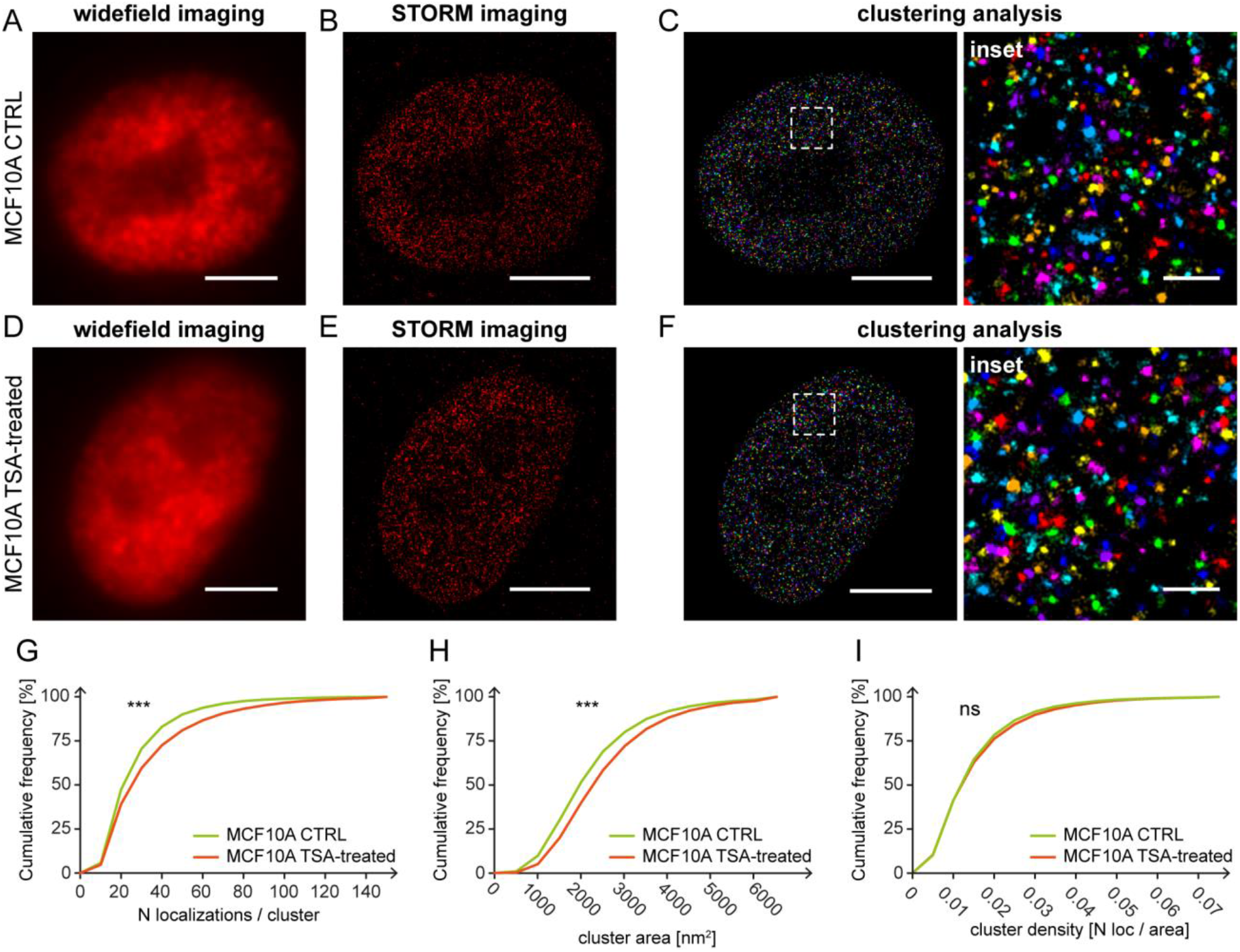
Super-resolution imaging of DEK protein clusters in control and TSA-treated MCF10A cells. (A) Widefield, (B) STORM images and (C) clustering image of DEK protein in control nucleus immunolabelled with AF647 (each color represents different cluster ID). (D) Widefield, (E) STORM images and (F) clustering image of DEK protein in TSA-treated nucleus immunolabelled with AF647 (each color represents different cluster ID). Cumulative distributions of (G) the number of localizations / cluster, (H) cluster area and (I) cluster density show the DEK cluster features differences between control and TSA-treated cells. The average number of localized events in control cells: 27.2 ± 18.7 compared to TSA-treated cells: 34.0 ± 26.3. The average cluster areas control cells: 2224.9 ± 1252.8 [nm^2^] compared to TSA-treated cells: 2543.1 ± 1369.0 [nm^2^]. The average cluster densities in control cells: 0.0147 ± 0.011 [N_loc_/cluster] and TSA-treated cells: 0.0153 ± 0.0118 [N_loc_/cluster]. The total number of analyzed clusters is: N_control_ = 26906 and N_TSA-treated_ = 21914, for n ≥ 3 independent experiments. Kolmogorov-Smirnov statistical test: not significant, n.s. when p > 0.05; “ *** “ when p ≤ 0.001. Scale bar = 5 µm. Scale bar insets = 500 nm.

**Fig. 4.**
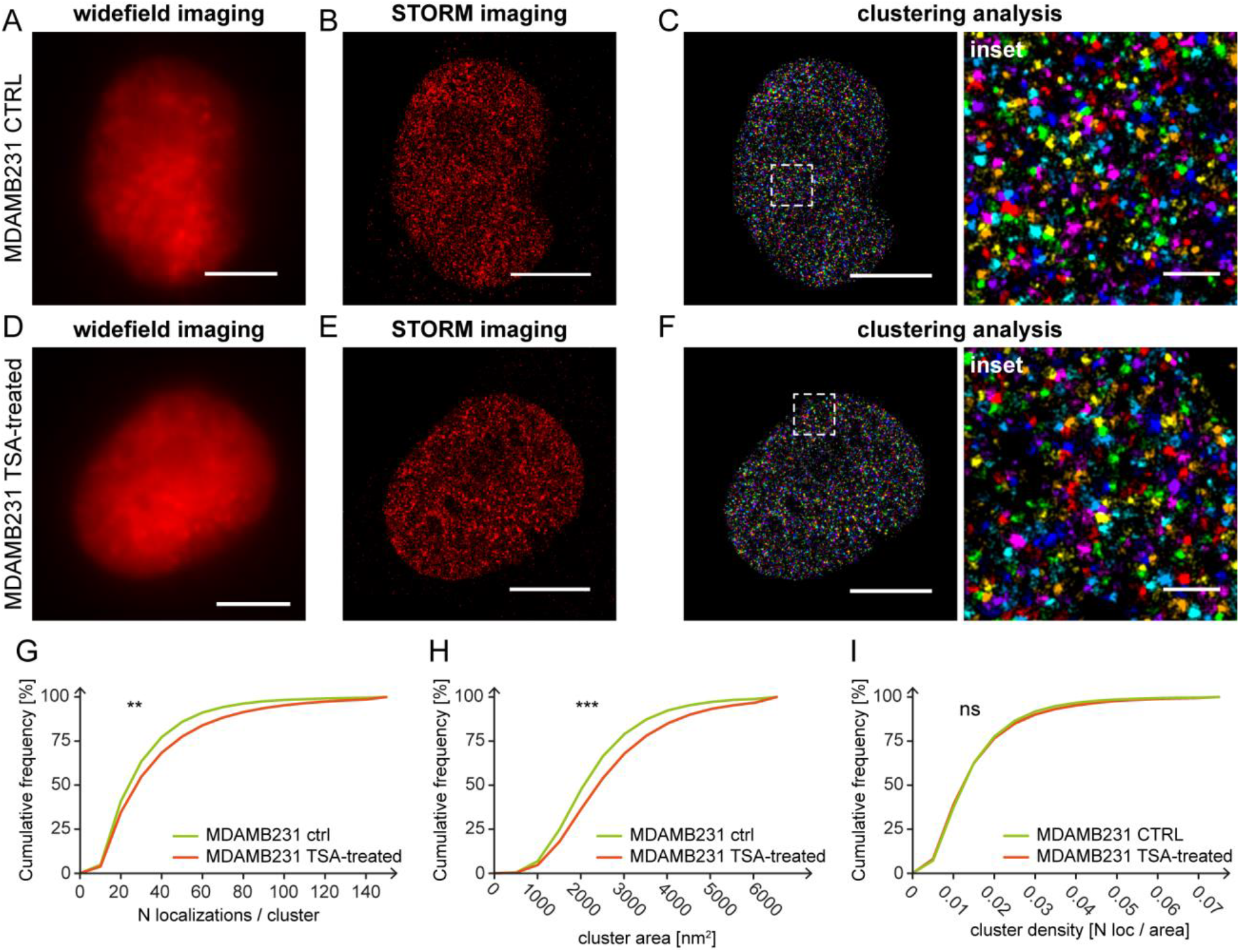
Super-resolution imaging of DEK protein clusters in control and TSA-treated MDAMB231 cells. (A) Widefield, (B) STORM images and (C) clustering image of DEK protein in control nucleus immunolabelled with AF647 (each color represents different cluster ID). (D) Widefield, (E) STORM images and (F) clustering image of DEK protein in TSA-treated nucleus immunolabelled with AF647 (each color represents different cluster ID). Cumulative distributions of (G) the number of localizations / cluster, (H) cluster area and (I) cluster density show the DEK cluster features differences between control and TSA-treated cells. The average number of localized events in control cells: 30.7 ± 21.9 compared to TSA-treated cells: 37.5 ± 29.9. The average cluster areas control cells: 2282.8 ± 1131.1 [nm^2^] compared to TSA-treated cells: 2687.3 ± 1450.7 [nm^2^]. The average cluster densities in control cells: 0.0151 ± 0.0106 [N_loc_/cluster] and TSA-treated cells: 0.0156 ± 0.0121 [N_loc_/cluster]. The total number of analyzed clusters is: N_control_ = 22008 and N_TSA-treated_ = 47585, for n ≥ 3 independent experiments. Kolmogorov-Smirnov statistical test: not significant, n.s. when p > 0.05; ; “ ** “ when p ≤ 0.01; “ *** “ when p ≤ 0.001. Scale bar = 5 µm. Scale bar insets = 500 nm.

The analysis of the distributions of the DEK clusters densities (Fig. 3 I) revealed that there were no statistically significant differences between control and TSA-treated cells for both cell lines. This result suggests that the local density of DEK within clusters is conserved not only regardless of the DEK expression level but also regardless of the chromatin compaction. In other words, despite the nanoscale rearrangement of DEK after TSA treatment demonstrated by the number of localizations per cluster and area differences, the overall density is maintained constant and this feature is conserved across cell types and conditions, suggesting the preservation of the DEK protein functioning.

## Discussion

It is well established that the DEK protein is overexpressed in highly proliferating cells (38), such as cancer cells, and can alter the chromatin topology (7). Our recent study using super-resolution techniques suggested that the commonly observed dispersed nuclear DEK pattern (18,31) is composed of nanoclusters (10). In this work, we investigated the nanoscale characteristics of such clusters in normal-like and aggressive type of breast cancer cell lines to study if DEK’s overexpression correlates with its cluster features. Moreover, we showed how the chromatin decompaction influences the DEK clusters in both cell lines.

Over 20 years ago, it has been shown that DEK stays in ratio of two to three DEK copies per nucleosome *in vitro* (7), and 0.25 – 0.5 DEK copy per nucleosome *in vivo* (4), depending on the proliferative state of the cell and their cell type. Interestingly, it has also been reported that DEK can multimerize (39).

Our results show that the number of localizations per cluster depends on the cell type, and more precisely could be related to the DEK expression level. MDAMB231 cells nuclei contain DEK clusters with higher localizations per cluster than MCF10A nuclei. Interestingly, even if in case of MDAMB231 cells the DEK clusters exhibit a larger mean area than in MCF10A cells, the difference of their distributions is not statistically significant. However, what we observed was the conservation of DEK protein density within clusters in both cell types.

We further showed that the DEK density within clusters is maintained also after the TSA treatment. In the context of breast cancer, TSA has been extensively studied as a molecular target for anticancer therapy (40–42). It has been previously shown that histone modifications are crucial for the regulation of the accessibility of chromatin to DNA binding factors (43), and in cancerous cells, local chromatin environment can be perturbed (44). In case of global histone acetylation, the opening of the chromatin enhances gene transcription and increases accessibility for nuclear proteins such as RNA Polymerase II and other transcription factors (45,46). Studied with super-resolution methods, TSA treatment results in lower number of localizations per nanodomain and lower nanodomain areas of H2B histone clusters in hyperacetylated human fibroblasts in comparison to non-treated cells (34). TSA induces chromatin opening (47) and a decrease of DNA density associated with nucleosome clutches (37). These conditions induce the formation of a large number of newly accessible sites in the genome. Here, in this work, we found that under the global chromatin changes inflicted by the TSA treatment of both cell lines, DEK protein reorganizes into bigger clusters with higher number of localizations (around 25% and 22% higher number of localizations for MCF10A and MDAMB231 cells, respectively, and around 14% and 18% higher cluster area for MCF10A and MDAMB231 cells, respectively). This finding nicely demonstrates that upon the histone hyperacetylation, and increased accessibility to the nucleosomal DNA, chromatin-associated proteins can reorganize into bigger nanostructures. This observation stays in line with other works in which a redistribution of nuclear organization of chromatin-associated proteins has been observed (for example heterochromatin protein 1 (HP1)) (48). Furthermore, single-molecule localization imaging demonstrated the association of RNA Polymerase II to nucleosomes clutches of smaller size and fewer number of nucleosomes, which correspond to the clutches similar to the ones generated after TSA treatment (34). It is likely that under the condition of the hyperacetylation of histone tails and DNA decompaction in chromatin, DEK protein accessibility is higher. Indeed, in both investigated cell lines, TSA treatment induced the formation of bigger clusters with higher number of localizations. Interestingly, although the area and number of localizations per clusters change upon the hyperacetylation conditions, the density of DEK (expressed as the number of localizations per area) stays conserved.

These semi-quantitative observations suggest that the nano-environment of DEK clusters is maintained and is not influenced by overexpression of the protein itself or by the change of the chromatin compaction.

