## Supplementary Figure 1 for "Super-resolution microscopy reveals the nanoscale organization of the DEK protein cancer biomarker"

### Supplementary Figures

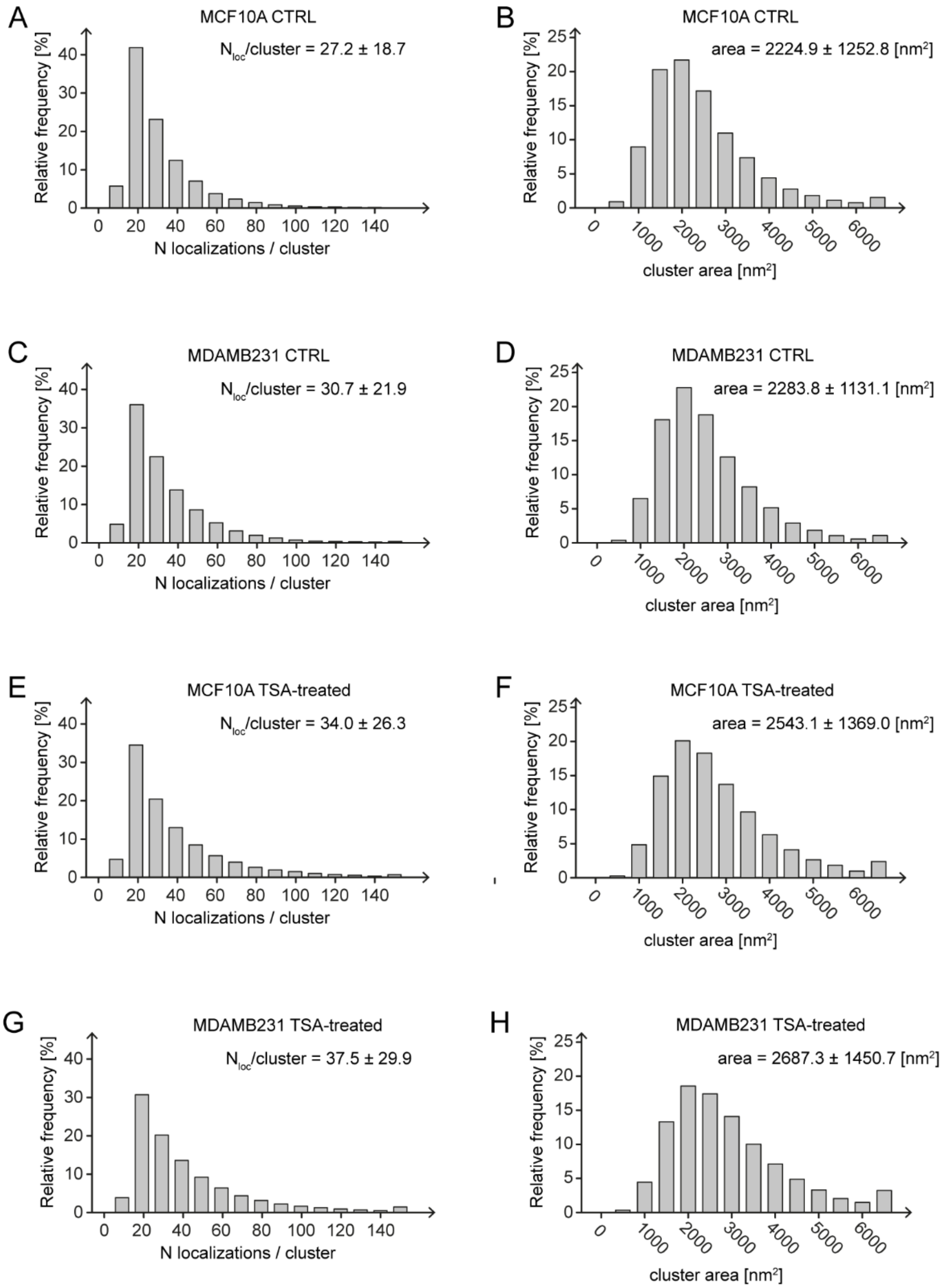


***Suppl. Fig. 1. Frequency distributions of number of localizations / cluster and cluster areas based on SMLM imaging.*** *The cluster parameters were measured in the immunolabelled with AF647 samples of: (A,B) MCF10A control cells, (C,D) MDAMB231 control cells, (E,F) MCF10A TSA-treated cells, and (G,H) MDAMB231 TSA-treated cells. The total number of analyzed clusters in MCF10A cells:* $N_{control}$ *= 26906 and* $N_{TSA\text{-}treated}$ *= 21914. The total number of analyzed clusters in MDAMB231 cells:* $N_{control}$ *= 22008 and* $N_{TSA\text{-}treated}$ *= 47585.*
